# Successive Bloodmeals Enhance Virus Dissemination Within Mosquitoes And Increase Transmission Potential

**DOI:** 10.1101/246306

**Authors:** Philip M. Armstrong, Hanna Ehrlich, Angela Bransfield, Joshua L. Warren, Virginia E. Pitzer, Doug E. Brackney

## Abstract

The recent Zika virus (ZIKV) and chikungunya virus (CHIKV) epidemics highlight the explosive nature of arthropod-borne (arbo) viruses transmitted by *Aedes aegypti* mosquitoes^1,2^. Vector competence and the extrinsic incubation period (EIP) are two key entomological parameters used to assess the public health risk posed by arboviruses^3^. These are typically measured empirically by offering mosquitoes an infectious bloodmeal and temporally sampling mosquitoes to determine infection and transmission status. This approach has been used for the better part of a century; however, it does not accurately capture the biology and behavior of many mosquito vectors which refeed frequently (every 2-3 days)^4^. Here we demonstrate that administration of a second non-infectious bloodmeal significantly shortens the EIP of ZIKV-infected *Ae. aegypti* by enhancing virus escape from the mosquito midgut. Similarly, a second bloodmeal increased the competence of this species for dengue virus and CHIKV. This effect was also observed for ZIKV in *Aedes albopictus,* suggesting that this species might be a more important vector than once thought and that this phenomenon may be common among other virus-vector pairings. Modeling of these findings reveals that a shortened EIP would result in a significant increase in the basic reproductive number, *R*_0_. This increase helps explain how *Ae.aegypti* can sustain an explosive epidemic like ZIKV despite its relatively poor vector competence in single-feed laboratory trials. Together, these data demonstrate a direct and unrecognized link between mosquito feeding behavior, EIP, and vector competence.

## MAIN TEXT

Arthropod-borne (arbo)viruses represent an ongoing threat to human health as shown by the emergence and global spread of dengue virus (DENV; *Flaviviridae*), chikungunya virus (CHIKV; *Togaviridae*), and Zika virus (ZIKV; *Flaviviridae*)^5,6^. These three arboviruses are transmitted by mosquitoes of the genus *Aedes* and are known to cause disease outbreaks with high attack rates, necessitating research into the factors regulating virus transmission. The urban-dwelling mosquito *Aedes aegypti* serves as a particularly efficient vector because it feeds predominately and frequently on human hosts (every 2-3 days) thereby increasing the frequency of host contact^7-9^. Nevertheless, in laboratory trials, *Ae. aegypti* populations from endemic regions often exhibit unexpectedly low vector competence values for their arboviruses as measured by the proportion of mosquitoes that become infected and transmit a pathogen after ingesting virus^10-14^. In these studies, mosquitoes are offered one initial bloodmeal, infected with the virus in question, and are not allowed to refeed again, as is standard practice for assessing vector competence. Therefore these studies do not recapitulate the natural biology of mosquitoes that refeed frequently. It is possible that differences in feeding history could help explain the seemingly paradoxical nature of *Ae. aegypti*-transmitted arboviruses.

Arboviruses must overcome multiple barriers to infection within the mosquito vector for transmission to occur^15^. The virus must infect the midgut, disseminate out of the midgut, traverse the basal lamina layer to the hemolymph, and then infect the salivary glands before being transmitted to the next vertebrate host^16^. Failure of virus escape out of the mosquito midgut to the peripheral tissues has been identified as an important barrier to arbovirus transmission, but the underlying factors mediating this process are poorly understood^10,11,17-19^. Blood feeding triggers physiological changes within the mosquito—including mechanical distention of the midgut, apoptosis and regeneration of midgut epithelial cells, and altered permeability of the basal lamina layer—that could enhance or accelerate virus dissemination out of the midgut^20,21^. Based on these considerations, we evaluated the hypothesis that virus-infected mosquitoes fed an additional non-infectious blood-meal will more effectively disseminate and transmit virus than mosquitoes fed only once.

To test this hypothesis, we provided *Ae. aegypti* a second non-infectious blood meal 3-4 days after the infectious blood meal and compared virus infection and dissemination rates to mosquitoes that received a single bloodmeal (Fig. 1a). Midgut infection prevalence of ZIKV was similar in the single- and double-feed groups, but the percentage of mosquitoes with disseminated ZIKV infection was significantly higher in mosquitoes receiving a second bloodmeal than those fed only once (Fig. 1b). Enhanced virus dissemination did not occur when mosquitoes were fed a non-infectious bloodmeal prior to receiving a ZIKV infectious bloodmeal (Fig. S1), suggesting that enhanced virus dissemination only occurs when virus is already seeded in the midgut prior to the non-infectious bloodmeal.

**Figure 1.**
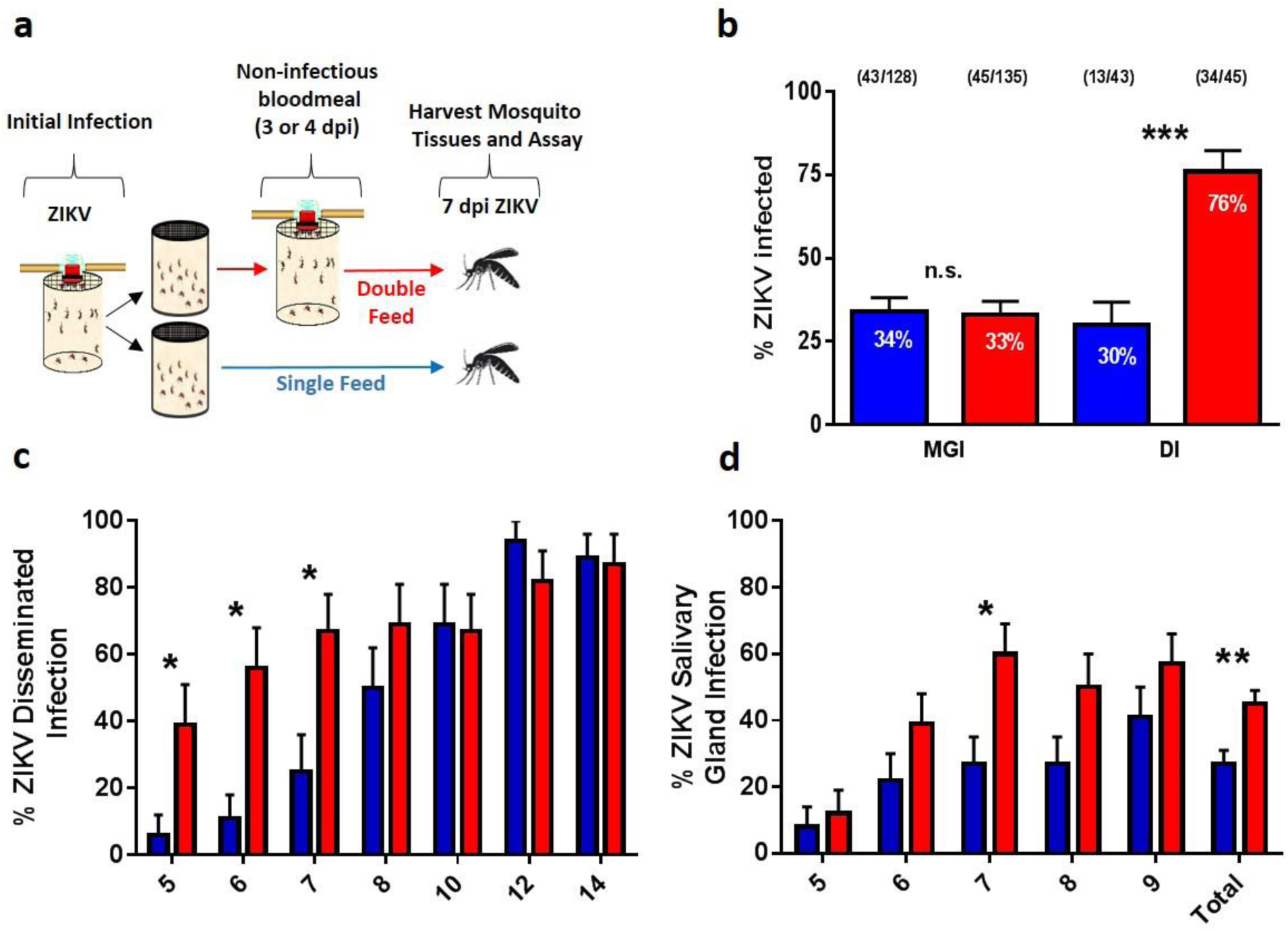
Multiple feeding events increase dissemination rates of ZIKV in *Aedes aegypti*. (a)Schematic of experimental design. *Aedes aegypti* mosquitoes were offered a ZIKV infectious bloodmeal and at three days post infection (dpi) individuals in the double-feed groups were fed a second, non-infectious bloodmeal. (b) At 7 dpi paired bodies and legs were collected and assayed for the presence of viral RNA. MGI (midgut infection rates; % of mosquitoes with viral RNA in their bodies), DI (disseminated infection; % of body positive mosquitoes with viral RNA in their legs). (c) Similarly, at 5-14 dpi paired bodies and legs were assayed for ZIKV RNA; only DI rates are presented (n=20/ treatment/ time point). (d) 5-9 dpi paired bodies and salivary glands were assayed for ZIKV RNA; only SGI rates are presented (n=34-39/ treatment/ time point). ‘Total’ represents the combined data from days 5-9 (n=183-187/ treatment). (●) single-feed; (●) double-feed. Data were analyzed by Fisher’s exact test. (*) p<0.05, (**) p<0.01, (***) p<0.001.

To assess virus dissemination dynamics and transmission potential, *Ae. aegypti* were fed ZIKV, refed a non-infectious bloodmeal 3 days later, and assayed for disseminated infection 5-14 days post-infection. ZIKV disseminated more rapidly in the double-feed group than the single-feed group, but the difference in dissemination rates disappeared by day 10 post-infection (Fig. 1c). The increase in early dissemination correlated with an increase of ZIKV found in the salivary glands and thus enhanced early transmission potential (Fig. 1d). These data demonstrate that the observed effects of double feeding shortened the extrinsic incubation period (EIP) of ZIKV in *Ae. aegypti* mosquitoes. To determine whether this was mediated by an influx of energy-rich blood promoting viral replication and midgut escape, we compared ZIKV titers in mosquito midguts (Fig. S2). The number of virus genome equivalents was similar in mosquitoes regardless of feeding status (single- versus double-feed) or infection status (midgut restricted versus disseminated infection). These findings are consistent with the literature, as others have demonstrated that there is no correlation between midgut titers and dissemination rates^22,23^, and indicate that once ZIKV has established an infection in the gut, its ability to escape is not conditioned by enhanced viral replication.

We subsequently tested ZIKV infection and dissemination in a low-generation (F5) colony of *Ae. albopictus* to determine whether a second bloodmeal increases vector competence of other mosquito species. Similar to *Ae. aegypti*, *Ae. albopictus* will refeed frequently in the field, taking multiple bloodmeals within a single gonadotrophic cycle^24^. Previous studies have shown that *Ae. albopictus* have low competency for ZIKV and as a result were thought to play a minor role during the ZIKV epidemic^25^; nevertheless, these studies did not consider the impact of refeeding on vector competence. Accordingly, *Ae. albopictus* were fed a ZIKV-infectious bloodmeal, offered a non-infectious bloodmeal 4 days later, and then tested for viral infection in mosquito bodies, legs, and salivary glands 7 and 10 days post infection (Fig. 2a). Administration of a second bloodmeal after infection significantly increased the percentage of mosquitoes with disseminated viral infections (Fig. 2b and 2c); this led to a higher percentage of salivary gland infections when expressed as the proportion of all ZIKV-exposed mosquitoes (Fig. 2d). Further, it was determined that the timing of the second bloodmeal in relation to the initial infectious bloodmeal did not influence the outcomes (Fig. S3). Unlike *Ae. aegypti*, increases in virus dissemination did not disappear by day 10 in *Ae. albopictus*. These findings suggest that under field conditions of frequent feeding, *Ae. albopictus* are more competent and could have contributed a larger role during the ZIKV epidemic than previously thought.

**Figure 2.**
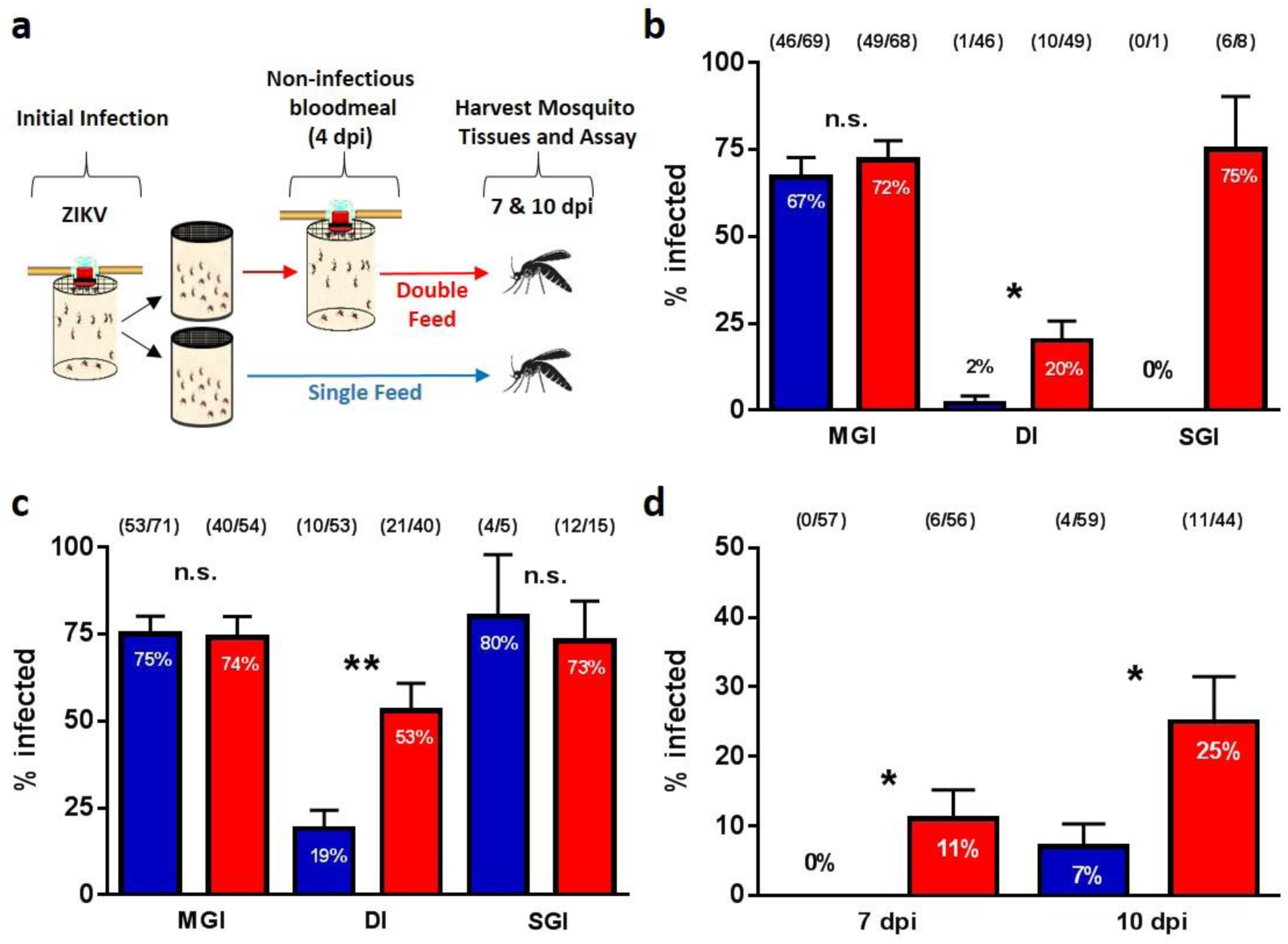
Multiple feeding events increase the potential of *Aedes albopictus* to transmit ZIKV. (a) Schematic of experimental design. *Aedes albopictus*mosquitoes were offered a ZIKVinfectious bloodmeal and four days post infection (dpi) individuals in the double-feed groups were fed a second, non-infectious bloodmeal. Paired bodies (MGI; midgut infection; % of mosquitoes with viral RNA in their bodies), legs (DI; disseminated infection; % of body positive mosquitoes with viral RNA in their legs) and salivary glands (SGI; salivary gland infection; % of leg positive mosquitoes with viral RNA in their salivary glands) were collected and assayed for the presence of viral RNA (b) 7 dpi and (c) 10 dpi. (d) SGI data from (a) and (b) analyzed as the of ZIKV-exposed mosquitoes with a salivary gland infection. The data presented represents at least three experimental replicates. (●) single-feed; (●) double-feed. Data were analyzed by Fisher’s exact test. (*) p<0.05, (**) p<0.01, (***) p<0.001.

To quantify how a second bloodmeal would affect transmission of ZIKV by *Ae. aegypti* as measured by the basic reproductive number (*R_0_*), we first combined our single-feed data with published data on ZIKV to estimate a mean EIP of 10.0 days (95% credible interval (CI): 5.1-20.3 days) with a standard deviation of 5.1 days (95% CI: 3.0-9.2 days) (Fig. 3a). Using our temporal double-feed data (Fig. 1c), the estimated mean EIP was 7.25 days with a standard deviation of 6.1 days (Fig. 3b). Fig. 3c displays the estimated EIP probability densities for mosquitoes offered one versus two bloodmeals based on the posterior mean parameter estimates. The posterior probability that the mean for the single-feed EIP (*μ_EIP__SF_*) was larger than that of the double-feed EIP (*μ_EIP__DF_*) is 0.913 (P(*μ_EIP__SF_* > | *μ_EIP__DF_ data*)= 0.913). Based on the traditional single-feed distribution of the EIP, we estimated the mean *R*_0*SF*_ (*μ_R0SF_*) to be 2.94 (95% CI: 1.80-4.28), whereas when mosquitoes were fed a second bloodmeal following the initial infectious bloodmeal, the mean *R*_0*DF*_ (*μ_R0DF_*) was 4.21 (95% CI: 3.26-5.61) (P(*μ_R0DF_* > *μ_R0SF_* | *data*) = 0.934) (Fig. 3d). The EIP was the first- or second-most influential parameter affecting the difference in *R*_0_ (*R*_0*DF*_ − *R*_0*SF*_) according to our two sensitivity analyses (Fig. S6). The estimated increase in *R*_0_ following a second bloodmeal may help explain the magnitude of *Ae. aegypti*-vectored ZIKV epidemics despite the relatively low competence observed experimentally (after a single infectious bloodmeal) for this vector.

**Figure 3:**
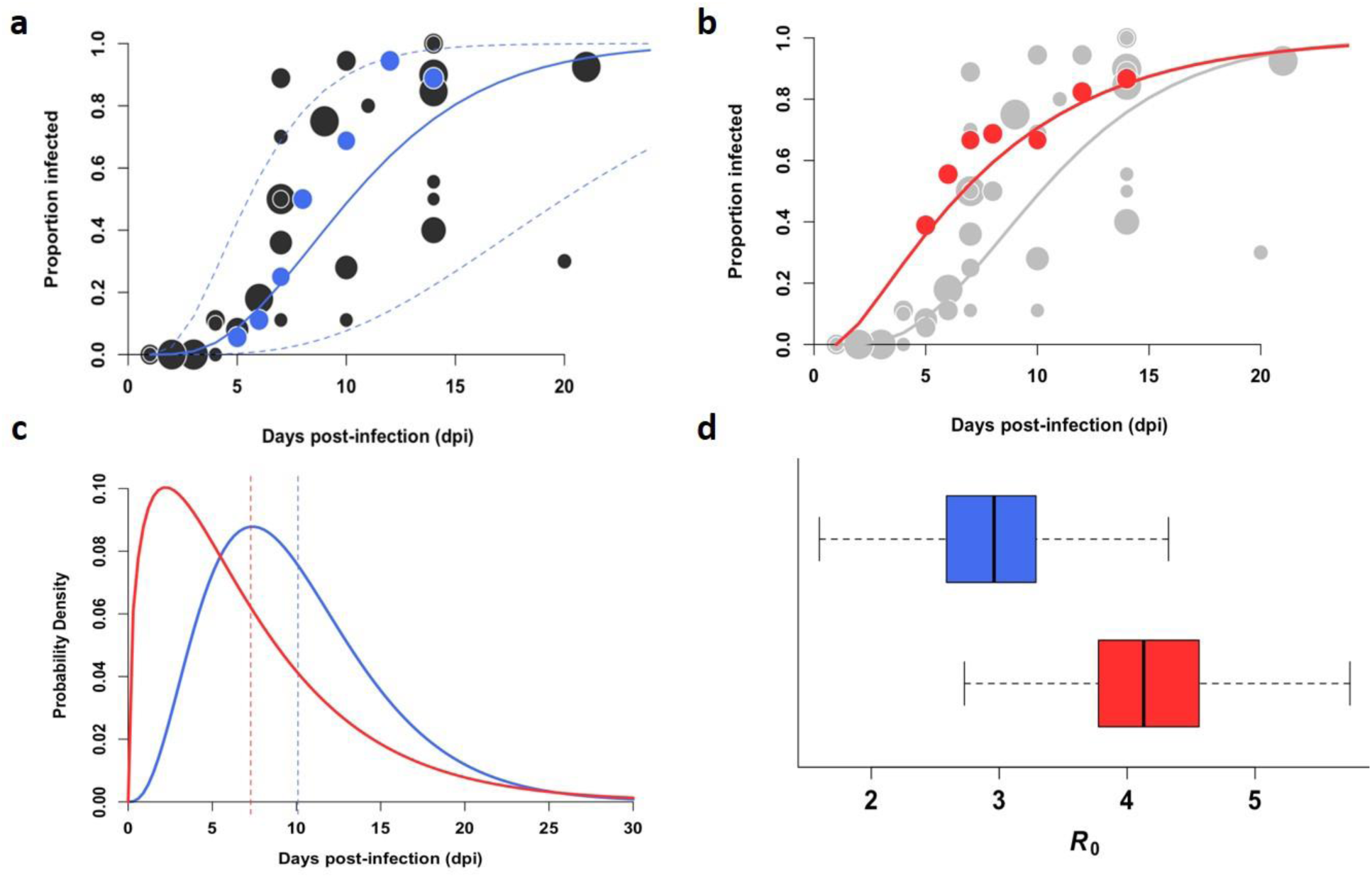
Offering mosquitoes a second bloodmeal post-infection decreases mean EIP and increases transmission potential of ZIKV. (a) We estimated ZIKV dissemination rates in *Aedes aegypti* mosquitoes based on data aggregated from seven published studies (black) and our experimental results (blue) for mosquitoes offered only one infectious bloodmeal. Each point represents the proportion of infected mosquitoes at a single time point in one particular study, weighted by sample size, where larger sample sizes have larger radii and more weight in estimating parameters. Gamma CDF model fit is represented by the solid line, while the 95% credible interval is depicted using dashed lines. (b) Data from our double-feed experiments (red circles) and fitted gamma CDF (red line) are plotted; the single-feed data are shown in grey for comparison. (c) Using our gamma distribution parameter estimates, we plotted probability densities for the single-feed EIP (blue line) and double-feed EIP (red line) with corresponding mean estimates for each EIP shown by the dashed lines. (d) The boxplots display the distribution of mean *R0* values for mosquitos offered a single bloodmeal (blue) versus a second bloodmeal (red).

To assess the impact of mosquito refeeding on vector competency for other arboviruses, *Aedes aegypti* were orally-exposed to dengue virus type 2 (DENV-2) and CHIKV and then given a second bloodmeal 3-4 days later. Refeeding *Ae. aegypti* a second time had no effect on midgut infection rates but significantly increased the proportion of mosquitoes with disseminated infections for both DENV-2 (Fig. 4b) and CHIKV (Fig. 4c). This demonstrates that the serial feeding behavior of *Ae. aegypti* enhances transmission potential of taxonomically diverse arboviruses and suggests that the mechanisms mediating this observation may be universally applicable to other virus-mosquito pairings.

This study establishes a connection between mosquito feeding behavior and viral development within the vector that has direct impacts on the transmissibility and epidemic risk of arboviruses. We found that providing a second non-infectious bloodmeal to mosquitoes enhances viral escape from the midgut for a number of different virus-vector pairings. Although the magnitude of difference between the single- and double-feed groups varied for each experimental replicate, enhancement itself was observed in every replicate (Table S1). The precise mechanism mediating viral escape from the midgut is poorly understood, but is a prerequisite for arbovirus transmission to occur^17^. The basal lamina layer surrounding the midgut epithelium represents a physical barrier that is impermeable to virus particles^26^, yet viruses are still able to escape from the gut. While a second non-infectious bloodmeal is not required for escape, a recent study showed that the integrity of the basal lamina was compromised 24-32 hours post-bloodmeal coincident with increased collagenase activity^21^. Future studies are needed to identify mechanisms mediating arbovirus escape from the midgut and how multiple bloodmeals enhance this event. Taken together, our findings emphasize the importance of considering feeding behavioral traits when performing vector competence studies. Past studies may underestimate the risks of arbovirus transmission by measuring vector competence after only one feeding on infected blood.

**Figure 4:**
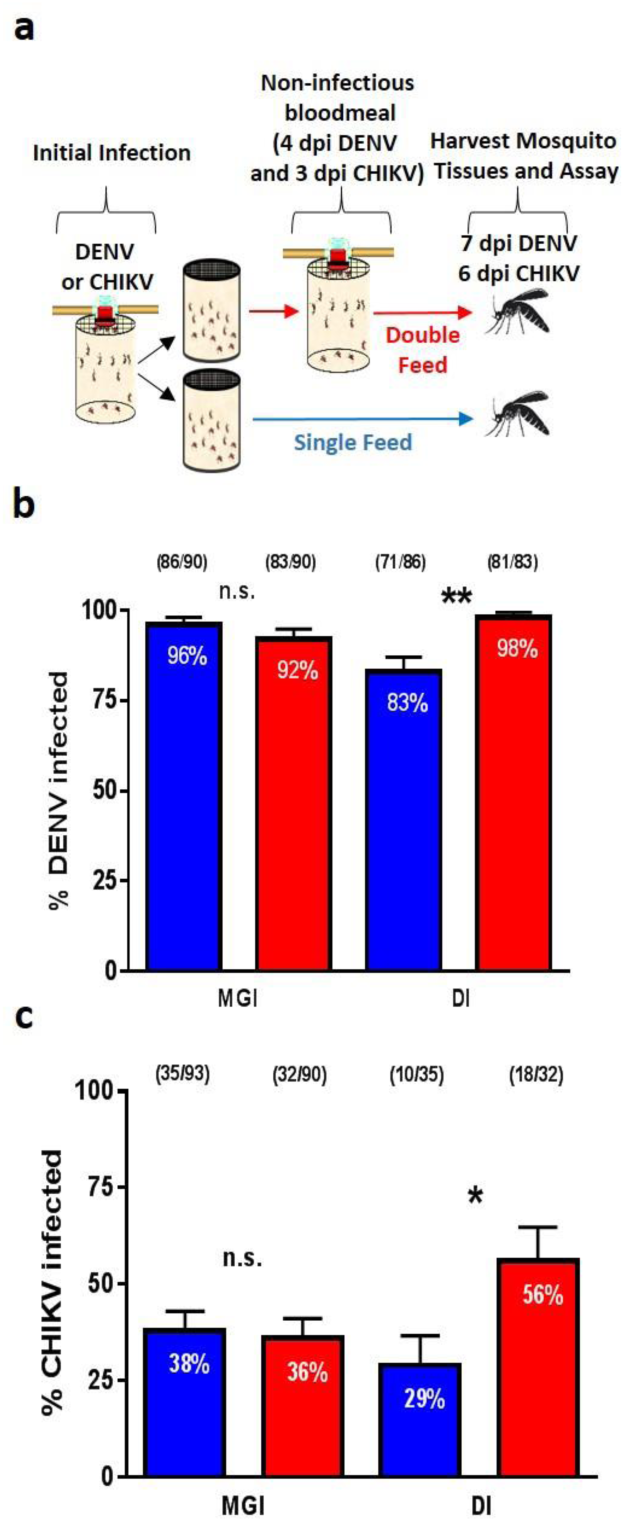
Multiple feeding events increase dissemination rates of DENV and CHIKV in *Aedes aegypti*. (a) Schematic of experimental design. *Aedes aegypti* mosquitoes were offeredeither a (b) DENV or (c) CHIKV infectious bloodmeal and at three (CHIKV) or four (DENV) days post infection (dpi) individuals in the double-feed groups were fed a second, non-infectious bloodmeal. At 6 dpi (CHIKV) and 7 dpi (DENV) paired bodies and legs were collected and assayed for the presence of viral RNA. MGI (midgut infection rates; % of mosquitoes with viral RNA in their bodies), DI (disseminated infection; % of body positive mosquitoes with viral RNA in their legs). The data presented represents two experimental replicates. (●) single-feed; (●) double-feed. Data were analyzed by Fisher’s exact test. (*) p<0.05, (**) p<0.01, (***) p<0.001.

## Acknowledgements

We thank John Shepard, Michael Misencik, and Maria Correa who provided technical assistance with rearing, processing, and virus testing of mosquitoes. This work was supported in part by grants from the Centers for Disease Control and Prevention (U50/CCU116806-01 and U01/CK000509-01), the US Department of Agriculture Hatch Funds (CONH00773), Multistate Research Project (NE1443), the National Center for Advancing Translational Sciences of the National Institutes of Health (CTSA grant number UL1 TR001863 and KL2 TR001862) (JLW), and the National Institutes of Health/National Institute of Allergy and Infectious Diseases (R01 AI112970) (VEP).

## METHODS

### Viruses, Cell Culture, and Mosquitoes

Viruses used in this study included ZIKV (PRVABC59; GenBank: KU501215), DENV-2 (125270/VENE93; GenBank: U91870), and CHIKV (R99659; GenBank: KX713902). C6/36 *Ae. albopictus* cells were used to amplify ZIKV and DENV-2, CHIKV were grown in Vero E6 cells, and BHK-21 (clone 15) cells were used to titrate infectious bloodmeals. Cell cultures were maintained in minimum essential medium (MEM) with 10% fetal bovine serum, 100 U/ ml penicillin, 100 mg/ ml streptomycin, 25 mg/ ml amphotericin B, L-glutamine, and sodium bicarbonate at 28°C for C6/36 cells or 37^o^ C for Vero and BHK-21 cells with 5% CO2. Colonies of *Ae. aegypti* (Orlando strain) and *Ae. albopictus* (Stratford strain, generation F5) were maintained on defibrinated sheep’s blood and reared under standard laboratory conditions. Adult mosquitoes were housed at 27°C in environmental chambers with a 14:10 light: dark cycle.

### Vector Competence Studies

*Ae. aegypti* and *Ae. albopictus* mosquitoes, 7-10 days post emergence, were offered an infectious bloodmeal containing a 1:1 mixture of defibrinated sheep’s blood and virus. After feeding, mosquitoes were cold-anesthetized and engorged females were transferred into two 32 oz. ice cream cartons containing a small cup with an egg-laying paper and housed in a 27°C environmental chamber. Three to seven days after the initial infectious bloodmeal, one of the two cartons was provided a second non-infectious bloodmeal. Again, engorged females were collected and placed in a carton with a new egg-laying cup and paper. Following variable extrinsic incubation periods (between 5-14 days post infection), mosquitoes were cold-anesthetized and bodies, midguts, legs and salivary glands were harvested. Forceps were flame sterilized between individual mosquito dissections, and salivary glands were washed twice in sterile PBS droplets prior to being immersed in 250 µl PBS-G [phosphate-buffered saline with 0.5% gelatin, 30% rabbit serum, and 1% 100x antibiotic-antimycotic (10,000 mg/ml of streptomycin, 10,000 U/ ml penicillin and 25 mg/ml of amphotericin B)] and macerated with a copper BB using a mixer mill. The inverse feeds (Fig. S1) followed a similar experimental design; however, the double-feed group received a non-infectious bloodmeal prior to receiving a ZIKV-infectious bloodmeal. Either freshly grown virus or frozen virus stocks were used to complete the studies. Initially, all of the ZIKV studies were performed with frozen stocks of ZIKV (4.8 x 10^6^ plaque forming units (pfu)/ml); however, in light of the poor midgut infection rates, all subsequent experiments were performed with freshly grown virus (1.0 x 10^6^ –10^7^ pfu/ml); C6/36 cells were infected at an multiplicity of infection of ~0.1 and harvested 4-5 days post infection. While this change in protocol did increase midgut infection rates, it did not alter the enhanced dissemination rate phenotype associated with multiple bloodmeals. DENV-2 was grown fresh on C6/36 cells (5 x 10^6^ – 3 x 10^7^ pfu/ml) and frozen aliquots of CHIKV (4 x 10^6^ pfu/ml) were used.

Infection rates were determined and reported as follows. Midgut infection (MGI) rate represents the total number of virus-positive bodies divided by the total number of mosquitoes tested. Similarly, disseminated infection (DI) rates were determined by dividing the total number of virus-positive legs by the total number of virus-positive bodies. Salivary gland infection (SGI) rates were determined by either dividing the number of virus-positive salivary glands by the total number of virus-positive bodies (Fig. 1d) or the total number of virus-positive legs (Fig. 2b and 2c) or the total number of blood-fed mosquitoes (Fig. 2d).

### Viral Detection by RT-qPCR

Total RNA was extracted from 50 µl of mosquito tissue and body homogenates using the Mag-Bind Viral DNA/RNA 96 Kit (Omega Bio-tek Inc., Norcross, GA) on a Kingfisher Flex automated nucleic acid extraction device (ThermoFisher Scientific, Waltham, MA) following the manufacturer’s instructions. Samples were eluted in 50 µl ddH2O. ZIKV RNA was detected in mosquito tissues using a previously described RT-qPCR primer-probe set (ZIKV 1087/1163c/1108 FAM)^27^. RNA standards were generated in order to quantify ZIKV RNA. Briefly, an ~ 680 bp fragment spanning the RT-qPCR primer set (positions 837-1520) was amplified with a forward primer containing a T7 promoter and a non-modified reverse primer. The amplicon was purified, sequenced, and used as template to generate RNA transcripts using the T7 Megascript Kit according to the manufacturer’s instructions (ThermoFisher Scientific). RNA was quantified on a Qubit Fluorometer (ThermoFisher Scientific) and diluted to achieve serial 10-fold genome equivalent (GE) dilutions. We detected 10^2^-10^7^ ZIKV GE/ reaction with a primer efficiency of 78.4% with an R^2^ value of 0.971, a slope of -3.977, and y-intercept = 46.965. DENV-2 RNA was detected using a previously described primer-probe set spanning the 3’-UTR^28^ and CHIKV RNA was detected using the previously described 6856F/6981c/6919-FAM primer-probe set^29^. The same RT-qPCR protocol was used to detect all three viruses. In brief, 25 µl reactions containing 2.5 µl of total RNA were assayed with the TaqMan RNA-to-Ct 1-Step Kit (ThermoFisher Scientific) on a CFX96 Touch Real-Time PCR Detection System (Bio-Rad, Hercules, CA) using the following parameters: RT - 50°C for 30 min, 95°C for 10 min, PCR - 95°C for 15 s., 60°C for 1 min followed by a plate read (50 cycles). Data were analyzed using the Bio-Rad CFX Manager 3.1 software and Ct values <35 cycles were considered positive for ZIKV and CHIKV and <33 cycles for DENV-2.

### Statistical Analysis of Experimental Data

The data were pooled from two to three independent replicates for each experiment involving mosquitoes (Table S1). Statistical methods were not used to predetermine sample size. Differences in the proportion of mosquitoes with midgut, disseminated, or salivary gland infection were analyzed using Fisher’s exact test. Standard error bars were determined by calculating the standard error of sample proportions. Descriptive statistics are provided in the figure legends. All analyses were performed using GraphPad Prism Statistical software.

### Estimation of the Extrinsic Incubation Period Distribution

In order to quantify how the additional bloodmeal would affect the transmission potential of ZIKV, as measured by the basic reproductive number *R*_0_, we first conducted a meta-analysis to characterize the EIP distribution of ZIKV. We identified ten studies that experimentally assessed the temporal profile of ZIKV dissemination in *Ae. aegypti* following a single infectious bloodmeal (Table S2). In each study, mosquitoes were infected orally with ZIKV and tested at varying time intervals to determine the proportion of infected mosquitoes at critical anatomical points (midgut, legs or wings, and salivary glands). We aggregated data on dissemination to the wings or legs (n=38 total observations from 8 studies, including the observations from our single feed experiment) and salivary gland infection (n=43 observations from 8 studies) separately, and then analyzed data from our double-feed experiments separately.

We used a Bayesian approach to estimate the EIP distribution for ZIKV. We assumed that the proportion of mosquitoes with disseminated infection by day *t* in experiment *i* follows a gamma cumulative distribution function (CDF)^30,31^, such that:

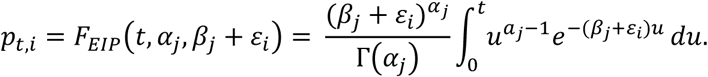

The gamma CDF is defined by shape parameter and rate parameter *βj* where *j*=1 for single-feed and *j*=2 for double-feed data. We included an additional parameter to account for between-study variability in the single-feed data. The observed number of mosquitoes with disseminated infection at time *t* in experiment *i* (*xt,i*) was assumed to be binomially distributed with sample size (*nt,i*) and a success probability (*pt,i*).

Posterior distributions of model parameters were estimated via a Markov chain Monte Carlo sampling algorithm implemented using JAGS, run from the statistical program R using the rjags package ^32^. We selected weakly informative prior distributions for the model parameters (see Supplementary Materials). The algorithm was run for 100,000 iterations with 5,000 burn-in iterations for two chains. Convergence was assessed through visual inspection of trace plots and calculation of the Gelman–Rubin convergence diagnostic (97.5% quantile 
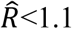
)
for all monitored parameters^33^.

We fit the model to both the DI and SGI data, but because of the similarity between the two distributions, we chose to utilize only the dissemination data to characterize the ZIKV EIP (Fig. S4) (Supplementary Materials). We then fit the same model—excluding the study variability parameter—to our double-feed data. To assess the difference in posterior distributions, we calculated the proportion of times that the mean EIP value (*¼_EIP__SF_*) for a thinned subset of posterior samples of the single-feed model was larger than that of the double-feed model (*¼_EIP__DF_*) (Fig. S5).

### Determining the Basic Reproductive Number (*R_0_*)

The basic reproduction number *R*_0_ is defined as the average number of secondary human cases that a primary human case generates over the course of infection in a fully susceptible population. For vector-borne diseases, *R*_0_ is typically estimated according to the Ross-MacDonald formula^34,35^:

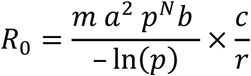

where *m* is the ratio of mosquito to human population density*, a* is rate at which mosquitoes bite humans, *p* is the probability of daily survival for mosquitoes, *b* is the vector competence, *N* is the EIP, *c* is the human infectiousness, and *r* is the human recovery rate. The biting rate *a* is expressed intrinsically as the product of the time interval between bloodmeals and the proportion of mosquito bloodmeals on humans. Traditionally, and for our single-feed *R*_0*SF*_, *a* is squared to account for the necessity of two bites to transmit virus from an infected to susceptible human. To calculate *R*0 for the double-feed data (*R*_0*DF*_), we modified this equation to examine the impact of an additional bloodmeal—which we assumed could come from either a human or non-human vertebrate—by multiplying by an additional factor *a*/*h*, where *h* is the proportion of bloodmeals taken from humans (see Supplementary Materials).

We parameterized our model according to current understanding of ZIKV transmission dynamics; parameter values and ranges are summarized in Table S3. We examined the impact of an additional bloodmeal by estimating the posterior distributions of mean *R*_0*SF*_ and *R*_0*DF*_ while drawing 10,000 random samples of *N* from the respective single- and double-feed EIP posterior distributions, holding all other parameters constant at their mean values. We calculated the posterior probability that the mean *R*_0*DF*0_ was larger than the mean *R*_0*SF*_ by calculating the proportion of times the mean *R*_0*DF*_ was greater than the mean *R*_0*SF*_ in our posterior samples. To assess the influence of the estimated variation in the EIP for the single- and double-feed models, we performed sensitivity analyses in which all parameters were varied independently according to their highest and lowest values, as well as their specified distributions, in order to determine the most influential parameters (Fig. S6). We also used Latin Hypercube Sampling to quantify overall uncertainty in *R*_0_ with and without the additional bloodmeal.

